# A Model-Based Cost-Effectiveness Analysis of an Exercise Program for Lung Cancer Survivors Following Curative-Intent Treatment

**DOI:** 10.1101/533281

**Authors:** Duc Ha, Jacqueline Kerr, Andrew L. Ries, Mark M. Fuster, Scott M. Lippman, James D. Murphy

## Abstract

**Rationale:** The Institute of Medicine emphasizes care in the post-treatment phase of the cancer survivorship continuum. Physical exercise has been shown to be effective in improving physical function and quality of life in cancer survivors; however, its cost-effectiveness in lung cancer survivors is not well established.

**Objective:** We performed a model-based cost-effectiveness analysis of an exercise intervention in lung cancer survivors following curative-intent treatment.

**Methods:** We constructed a Markov model to simulate the impact of the Lifestyle Interventions and Independence for Elders (LIFE) exercise intervention compared to usual care for stage I-IIIA lung cancer survivors. Costs and utility benefit of exercise were extracted from the LIFE study. Baseline utilities, transition probabilities, and survival were modeled. We calculated and considered incremental cost-effectiveness ratios (ICERs) <$100,000/quality-adjusted life-year (QALY) as cost-effective, and assessed model uncertainty using one-way and probabilistic sensitivity analyses.

**Results:** Our base-case model found that the LIFE exercise program would increase overall cost by $4,740 and effectiveness by 0.06 QALYs compared to usual care, and have an ICER of $79,504/QALY. The model was most sensitive to the cost of the exercise program, probability of increasing exercise, and utility benefit related to exercise. At a willingness-to-pay threshold of $100,000/QALY, the LIFE exercise program had a 71% probability of being cost-effective compared to 27% for usual care. When we included opportunity costs, the LIFE exercise program had an ICER of $179,774/QALY, exceeding the cost-effectiveness threshold.

**Conclusions:** A simulation of the LIFE exercise program in lung cancer survivors following curative-intent treatment demonstrates cost-effectiveness from an organization but not societal perspective. Strategies to effectively increase exercise remotely may be more cost-effective than in-facility strategies for these patients.

## Introduction

Up to 50% of patients with non-small cell lung cancer (NSCLC) are diagnosed with stage I-IIIA disease,^1^ the treatment of which typical aims to cure through a combination of lung cancer resection surgery, definitive radiation, or concurrent chemoradiation. Following curative-intent treatment, lung cancer survivors are at risk for health impairments resulting from the effects and/or treatment of lung cancer and comorbidities. Perioperative pulmonary^2^ and cardiopulmonary^3^ complications occur in 15% and 35%, respectively of patients undergoing surgical resection and can lead to negative health consequences well beyond the perioperative period.^4^ Additionally, patients typically lose 10-15% of lung function at three^5^ to six^6^ months following lobectomy which can negatively affect exercise capacity and health. Chemotherapy and radiotherapy, often part of the treatment for stage IB-IIIA NSCLC, can also lead to long-term health impairments.^7^

Lung cancer patients also have major comorbidities^8^ including smoking-related chronic obstructive pulmonary disease (COPD, 53%), diabetes (16%), and heart failure (13%) which may limit health. In time, partly due to the long-term effects of lung cancer treatment and comorbidities, patients experience disabling symptoms and a downward spiral of health.^9^ As such, clinical guidelines recommend health monitoring and counseling for wellness/health promotion for cancer including lung cancer survivorship care.^10^

Physical activity and exercise has been shown to improve physical function and quality of life (QoL) in cancer survivors.^11^ As such, the American College of Sports Medicine recommends exercise training as an adjunct cancer therapy^12^ and the American Cancer Society recommends cancer survivors to participate in physical activity/exercise regularly.^13^ However, the cost-effectiveness of exercise in cancer survivors including lung is relatively unknown. Model-based cost-effectiveness analyses (CEAs) of exercise interventions to improve lung cancer outcomes may motivate future clinical trials to test the effectiveness and trial-based CEAs of exercise interventions in lung cancer survivors. In this study, we simulated an exercise program in lung cancer survivors following curative-intent treatment to evaluate its cost-effectiveness. We hypothesize that this exercise program will be cost-effective in improving QoL.

## Methods

### Study Overview

We developed a simulation model to investigate the cost-effectiveness of exercise interventions compared to usual care (UC) in stage I-IIIA lung cancer survivors following curative-intent treatment. We chose the interventions described in the Lifestyle Interventions and Independence for Elders (LIFE) study,^14^ partly due to a higher median age (70) at diagnosis for lung cancer patients^15^ compared to other cancers and the availability of a previous CEA.^16^ In LIFE, 1,635 participants aged 70-89 years were randomized to a structured moderate-intensity exercise program (done in eight different centers and at home) or a health education program; participants in the exercise program had a ∼20% lower risk of major mobility disability compared to health education after a median of 2.6 years of follow-up.^14^ We created three different intervention conditions: participation in the LIFE exercise program, health education, or UC (Figure 1). Our outcome was the incremental cost-effectiveness ratio (ICER), expressed in US dollars per quality-adjusted life-years ($/QALYs).

**Figure 1:**
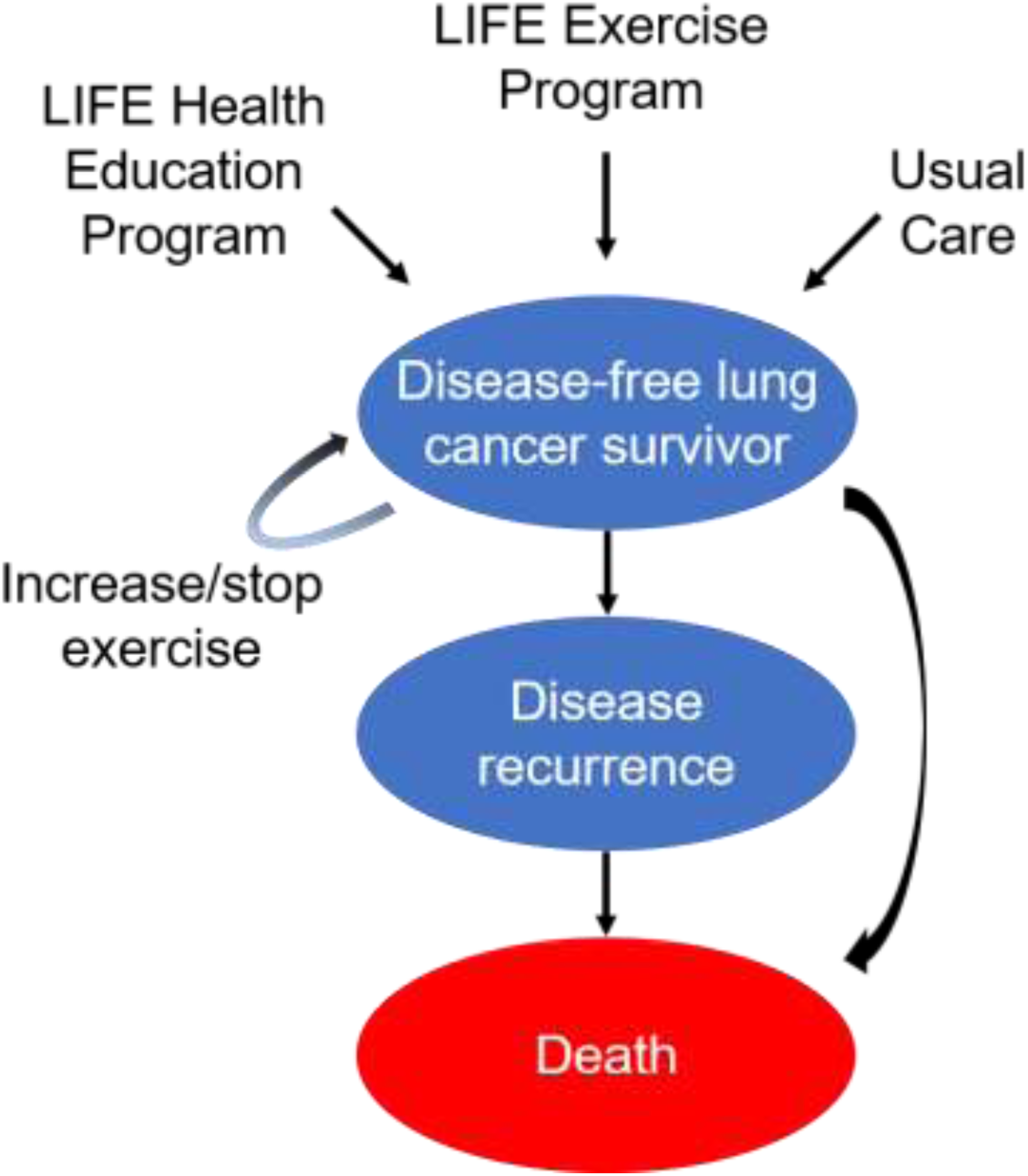
State Transition Diagram Caption: The health states are represented by ovals and include “disease-free, increase exercise,” “disease-free, no change in exercise,” “disease recurrence,” and “death.” Arrows represent possible transitions from one health state to another. Patients in the “disease-free” health state may experience a health utility benefit related to exercise if they increase their physical activity/exercise. Patients in the “disease recurrence” health state are assumed to not exercise. Patients who increase their exercise are subjected to a chance of stopping exercise after each year. LIFE = Lifestyle Interventions and Independence for Elders study

### Utilities

We used baseline health utilities as assessed by the EuroQoL-5 Dimensions (EQ-5D) questionnaire and reported in the Cancer Care Outcomes Research and Surveillance Consortium for lung cancer survivors who completed treatment >30 days previously:^17^ 0.83 for stage I patients, 0.78 for stage II, 0.76 for stage III, calculated to an average (0.79) for all stage I-IIIA patients based on estimates of stage distribution by the American College of Chest Physicians. For the disease recurrence health state, we assumed a health utility equivalent to stage IV patients (0.76) (Table 1).^17^

**Table 1:** Parameters for Cost-Effectiveness Model

| Variable | Base Case | Citation | Range Tested | Distribution |
| --- | --- | --- | --- | --- |
| Age | 70 | SEER<sup>15</sup> | 55 – 85 | Binomial |
| Five-year disease-free survival | 0.61 | Dziedzic<sup>23</sup> | 0.25 – 0.8 | Beta |
| Five-year overall survival | 0.55 | SEER<sup>15</sup> | 0.3 – 0.6 | Beta |
| Health utilities |  |  |  | Beta |
| Disease-free | 0.79 | Tramontano<sup>17</sup> | 0.76 – 0.83 | Beta |
| Disease-recurrence | 0.76 | Tramontano<sup>17</sup> | 0.70 – 0.80 |  |
| Benefit related to exercise* | 0.05 | Groessl<sup>16</sup> | 0.01 – 0.10 |  |
| Costs (\$/patient/year) | | | | Gamma |
| Exercise interventions** |  |  |  | Gamma |
| LIFE physical activity | 1,361 (1,855) | Groessl<sup>16</sup> | 500 – 2,500 |  |
| LIFE health education | 413 (1,210) | Groessl<sup>16</sup> | 250 – 1,000 |  |
| Usual care | 0 (203) | N/A<sup>†</sup> |  |  |
| Treatment of recurrent disease | 95,363 | Bradley<sup>20</sup> | 80,000-120,000 |  |
| Probability of increasing exercise |  |  |  | Beta |
| Physical activity/exercise | 0.60 | Gordon<sup>18</sup> | 0.40 – 0.80 | Beta |
| Health education | 0.10 | N/A<sup>†</sup> | 0.10 – 0.20 |  |
| Usual care | 0 | N/A<sup>†</sup> | 0 – 0.10 |  |
| Probability of dropout/lost to follow-up* | 0.10 | Speck<sup>19</sup> | 0.10 – 0.20 | Beta |

\*Parameter values at one year.
\*\*Opportunity costs entered in parentheses.
†Parameter assumed.
LIFE = Lifestyle Interventions and Independence for Elders study; SEER = Surveillance, Epidemiology, and End Results program

We assumed a health utility benefit related to exercise of 0.05 QALYs (as described in the LIFE study^16^) for patients in the disease-free and increase exercise health state. For patients in the LIFE exercise program, we conservatively estimated a probability of increasing exercise of 60% based on a pragmatic trial of breast cancer survivors^18^ and assumed probabilities of increasing exercise of 10% and 0% for those in the health education program and UC, respectively. We chose the pragmatic trial^18^ of breast cancer survivors due to previous evidence of effective exercise trials following treatment^19^ and the availability of a probability of increasing exercise.^18^ Patients who increase their exercise as the result of participating in the LIFE exercise or health education program were subjected to a yearly dropout rate of 10% based on a systemic review of controlled physical activity trials in cancer survivors.^19^

### Costs

We used costs as previously reported from an organizational perspective for the LIFE interventions^16^ and assumed a cost of US$0 for UC. We used the treatment cost for advanced-stage NSCLC as the cost for recurrent disease which included costs for targeted therapy but not immunotherapy.^20^ We also estimated the costs from a societal perspective as suggested by recommendations for the conduct, methodological practices, and reporting of cost-effectiveness analyses,^21^ and included participant opportunity costs related to time spent in exercise as reported in LIFE (213 min/week and 173 min/week in the exercise and health education programs, respectively)^14^ and assumed 35 min/week of exercise for those in UC based on data from the National Health and Nutritional Examination Survey for participants aged >70 years.^22^ We also included traveling costs for in-facility sessions using estimates of 40-min and 16-mile roundtrip for each session and a $2.42/gallon rate for gas cost and average light-duty vehicle efficiency of 23.9 miles/gallon. We measured the cost of time spent from lost work using the federal minimum wage ($7.25/hour as reported by the US Department of Labor). Given that the median age at diagnosis is 70 for lung cancer,^15^ we assume a substantial fraction of lung cancer patients would be retired, and assumed the lost opportunity costs of retired individuals were the same as the lost wages of younger individuals. We adjusted all costs for inflation using the Consumer Price Index calculator to estimate costs in 2017 (Table 1).

### Survival Probabilities

We obtained disease-free survival (DFS) data from the largest series to date^23^ involving 14,578 patients postresection for NSCLC and calculated a weighted 5-year DFS rate (61%) for patients aged 65-90 years, and 5-year overall survival (OS, 55%) from the Surveillance, Epidemiology, and End Results database^15^ for cases with segmentectomy or greater resection. We converted all 5-year to quarterly rates and calibrated them accordingly. We also incorporated the probability of dying of natural causes for a given age using data from the Social Security Actuarial Life Table.^24^

### Model Structure

We used TreeAge^®^ software to build a decision tree and capture outcomes including survival as described above, and Markov models to simulate health state progressions. Patients enter each intervention condition in equal probability and thereafter Markov models with four different health states: disease-free and increased exercise, disease-free with no change in exercise, recurrent disease, or death. Patients in the disease-free health state are subjected to a chance of dying of a natural death, dropout/stopping exercise if they increased exercise, and disease recurrence. Patients with disease recurrence will either continue or be subjected to death from lung cancer recurrence (Figure 1).

### Statistical Analysis

We converted all yearly to quarterly (1/4^th^) costs and utilities and used a cycle length of three months and time horizon five years with half-cycle corrections. We incorporated 3% annual discounts to costs and utilities,^21^ assumed a willingness-to-pay (WTP) threshold of $100,000 per QALY gained,^25^ and analyzed from healthcare organization and societal perspectives.^21^ We conducted sensitivity analyses on cost, probability of increasing exercise, and health utility benefit related to exercise to observe their effects on the ICER and probabilistic sensitivity analyses using Monte Carlo simulation with 100,000 iterations and parameters as reported in Table 1, with standard deviations 20% for all included variables.

## Results

The results of our model validation are shown in Figure 2.

**Figure 2:**
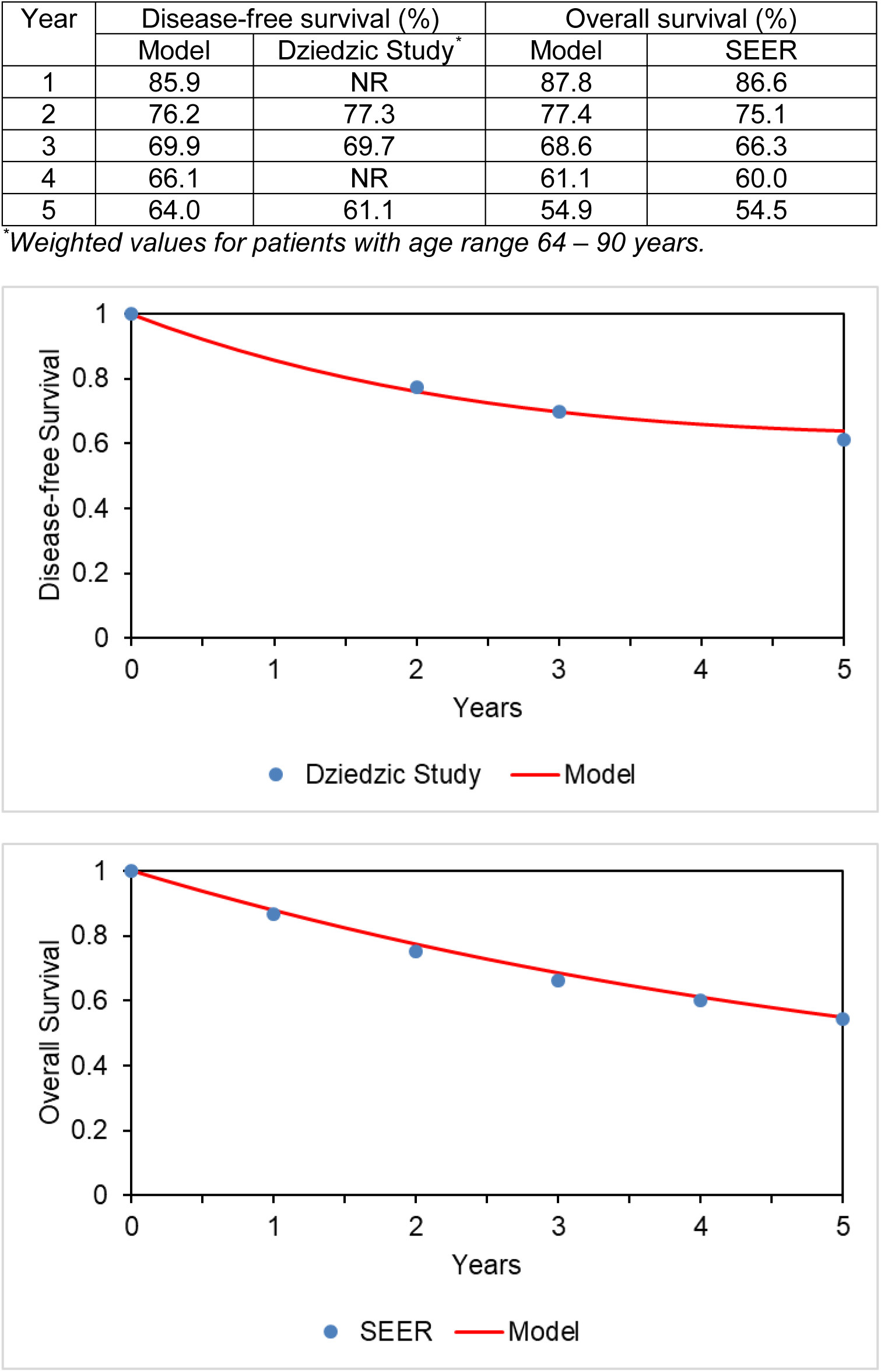
Validation of Model Endpoints Caption: Validation of model endpoints: (top) disease-free survival and (bottom) overall survival. Red colored lines represent our model data and blue dots represent comparison study/data. NR = not reported; SEER = Surveillance, Epidemiology, and End Results program

### Base Case Analysis

Our base case CEA from an organizational perspective found total costs of $110,224, $106,923, $105,485 and effectiveness 2.78, 2.73, and 2.72 QALYs for the LIFE exercise program, health education, and UC, respectively. The ICER for the health education program was higher than that of the LIFE exercise program, suggesting lack of cost-effectiveness (due to extended dominance). The incremental cost for the LIFE exercise program was $4,740 and incremental effectiveness 0.06 QALYs more than UC, giving an ICER of $79,504/QALY. The LIFE exercise program therefore would be cost-effective in lung cancer survivors from an organizational perspective at a WTP of $100,000/QALY. When we incorporated opportunity and traveling costs, the health education was again less cost-effective than the LIFE exercise program. The total cost for the LIFE exercise program was $116,685 and UC $105,967, giving an incremental cost of $10,718 and ICER $179,774/QALY. The LIFE exercise program therefore would not be cost-effective from a societal perspective when costs related to time lost to exercise and traveling were included. When we excluded traveling costs ($620/patient/year) to simulate an equally effective home-based intervention, the ICER was $143,556/QALY.

### One-Way Sensitivity Analyses

Our model was most sensitive to the costs of the exercise program, probability of increasing exercise, and health utility benefit related to exercise (Figure 3A-C). At a WTP threshold of $100,000, the cost of the exercise program could be up to a maximum of $1,712/patient/year (∼26% increase from our base case) and still be considered cost-effective compared to UC; a minimal probability of 48% of increasing exercise and utility benefit of 0.040 QALYs are needed to achieve cost-effectiveness.

**Figure 3:**
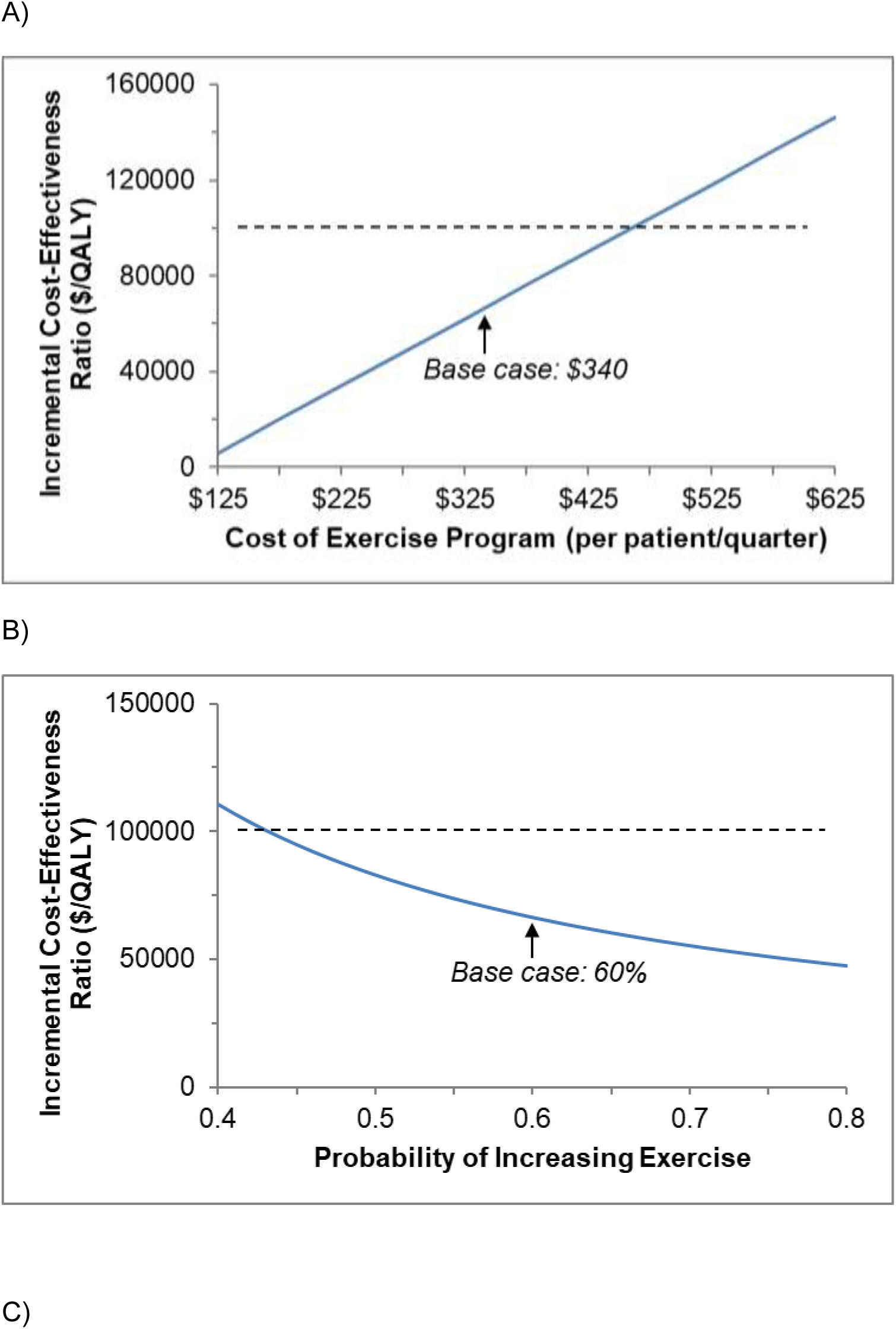
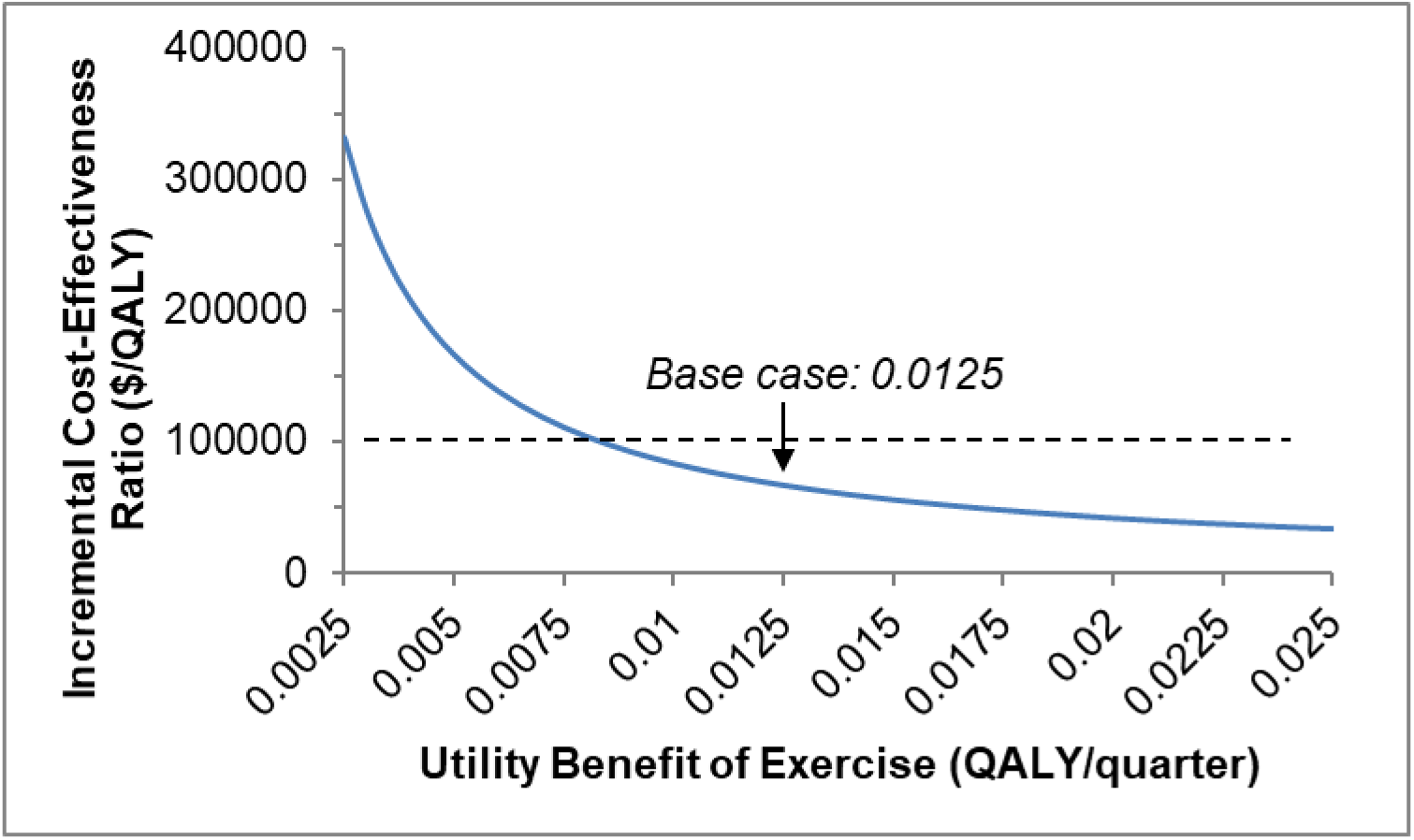
One-way Sensitivity Analyses A) Caption: Results of one-way sensitivity analyses for: (A) cost of exercise program, (B) probability of increasing exercise, and (C) utility benefit related to exercise. Dashed lines represent a willingness to pay threshold of $100,000, below which exercise would be considered cost-effective. QALY = quality-adjusted life-year

### Probabilistic Sensitivity Analyses

We conducted probabilistic sensitivity analyses by varying all costs, utilities, and probabilities simultaneously. At a WTP of $100,000/QALY, the LIFE exercise program was cost-effective 71% and UC 27% of the time (Figure 4). If we increased the WTP to $150,000,^25^ these respective probabilities were 94% and 4%.

**Figure 4:**
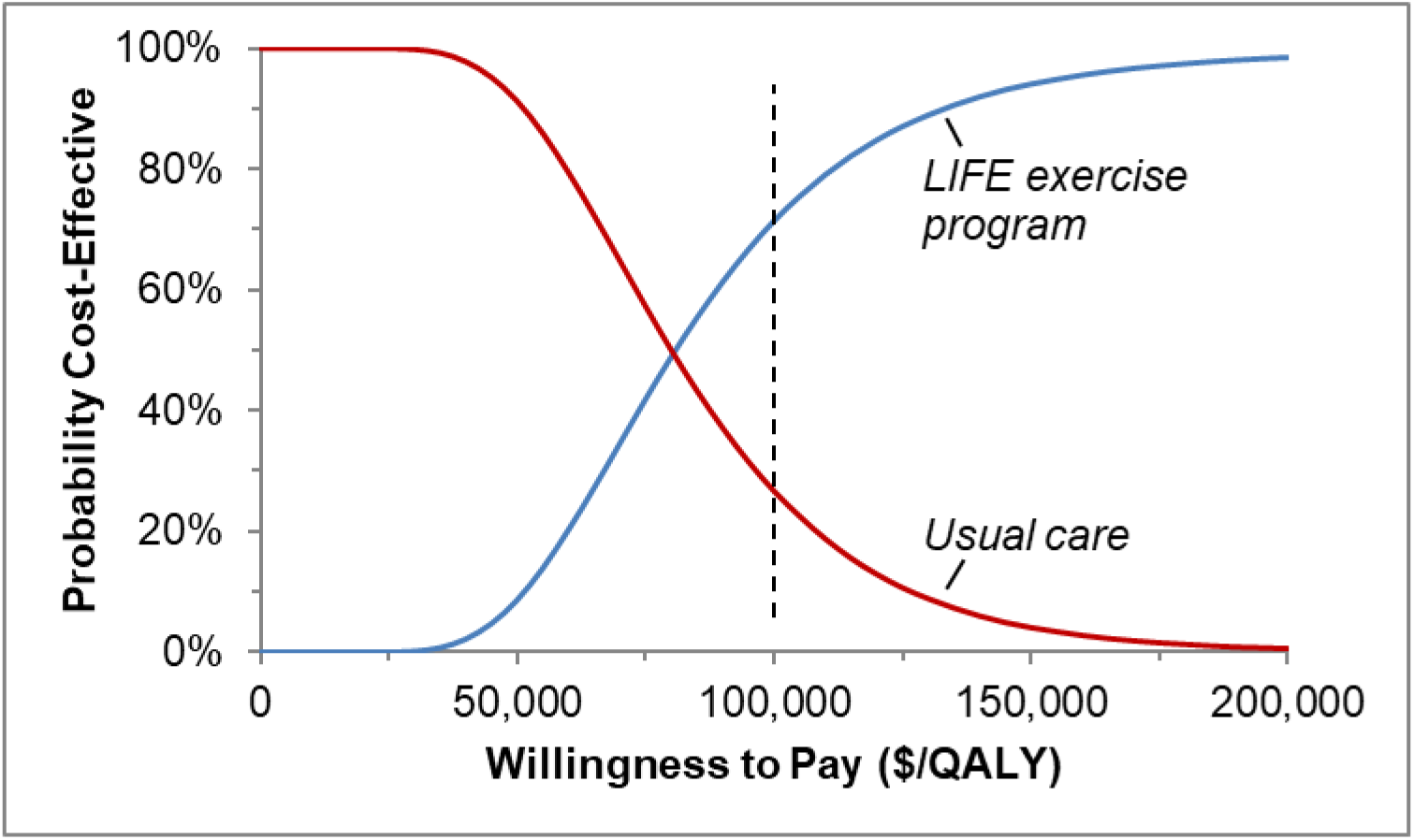
Cost-Effectiveness Acceptability Curve Caption: Results of probabilistic sensitivity analysis comparing the cost-effectiveness of the LIFE exercise program (blue line) with usual care (red line). Dashed line represents a willingness to pay threshold of $100,000/QALY, at which the probabilities of cost-effectiveness are 27% for usual care and 71% for the LIFE exercise program. LIFE = Lifestyle Interventions and Independence for Elders study; QALY = quality-adjusted life-year

## Discussion

In a model-based CEA, we found that the LIFE exercise program was cost-effective in lung cancer survivors following curative-intent treatment from an organizational but not societal perspective. These findings may have implications in program design and implementation in future studies aimed at improving exercise and QoL in these patients.

Our model is most similar to a previous analysis by van Waart and coworkers^26^ who performed a trial-based CEA of a home-based, low-intensity (Onco-Move) and a supervised, moderate-to-high intensity (OnTrack) program compared to UC. In 230 breast cancer patients enrolled from the initiation of adjuvant chemotherapy, Onco-Move was not cost-effective compared to OnTrack at ∼9.5 months. Compared to UC, OnTrack resulted in 0.04 QALYs gained and had ICERs of €21,721/QALY from a healthcare perspective and €26,916/QALY from a societal perspective (additionally included costs from informal care, absenteeism from work, and unpaid productivity losses). At a Dutch societal WTP of €80,000/QALY, OnTrack had a probability of cost-effectiveness of 79%. Interestingly, the authors found no significant differences in primary (e.g. general practitioner, allied health professions) or secondary (e.g. hospitalizations, outpatient clinics) healthcare costs between those in the OnTrack program compared to UC, but lower primary and higher secondary healthcare costs and no difference in total costs for Onco-Move compared to UC.

Similarly, Kampshoff and coworkers^27^ evaluated the long-term cost-effectiveness of a high-intensity (HI) compared to low-moderate intensity (LMI) exercise program at 64 weeks, and found that in 277 (65% breast and 18% colon) cancer survivors randomized to a 12-week HI or LMI program, followed by booster sessions at 4, 10, and 18 weeks, the HI program gained 0.03 QALYs and had ICERs of -€72,859/QALY (i.e. cheaper and more effective) and -€87,831/QALY compared to LMI from the healthcare and societal perspectives, respectively. Notably, the cost saving was primarily due to significantly lower healthcare costs in secondary care and medications for participants in the HI compared to the LMI program.

In contrast, May and coworkers^28^ performed a CEA for an 18-week exercise program during adjuvant breast and colon cancer treatment, and found that at 9 months, supervised exercise had incremental effectiveness of 0.01 and 0.03 QALYs for breast (n=204) and colon (n=33) cancer, respectively, and societal costs of €2,912 and -€4,321 compared to UC. Due to ICERs for breast cancer patients exceeding the WTP threshold, the authors concluded that the 18-week exercise program was cost-effective only in colon cancer, partly due to cost saving in chemotherapy, hospitalization, medical contacts, and work absenteeism for the exercise group compared to UC.

Like these CEAs, our study suggests that the cost-effectiveness of exercise programs largely depends on the effectiveness (QALYs) gained. While our model is also sensitive to the costs of the program, these trial-based CEAs alternatively suggest that exercise may be associated with lower healthcare costs and therefore cost-effective. Unlike these CEAs, our study also incorporated a probability of increasing exercise which indirectly estimates the probability of exercise uptake and therefore may have implications for generalization/translation in healthcare delivery. In addition, we estimated the traveling costs for facility-based exercise programs which may be associated with more QALYs gained^11^ but higher participant costs compared to home-based interventions. Our model was somewhat sensitive to the 5-year DFS but not OS rate, likely due to the assumptions that patients with disease recurrence do not exercise and there is no survival benefit related to exercise.

The estimated direct medical costs for cancer in the US was $80.2 billion in 2015, much of which (∼90%) was spent on hospital outpatient/office visits or inpatient hospital stays.^29^ Often overlooked is the economic impact of cancer survivorship in years after diagnosis. Based on the Medical Expenditure Panel Survey, the mean health expenditure and estimated annual lost productivity for cancer survivors is higher beyond one year of diagnosis compared to those without a history of cancer.^30^ In addition, cancer survivors are also more likely to have chronic conditions/multiple chronic conditions and therefore increased expenditure and productivity loss.^30^ As such, cancer survivors commonly report financial hardships and need access to lifelong personalized screening, surveillance, and chronic disease management.^30^

Following a cancer diagnosis, the Institute of Medicine recommends care planning, psychological support, and prevention and management of long-term and late treatment effects.^31^ The care plan for each cancer survivor should include access to supporting services and tips on maintaining a healthy lifestyle.^31^ While adult cancer survivors experience both physical and psychological impairments, evidence suggests that more cancer survivors have reduced QoL as the result of physical than psychological impairments;^32^ these impairments often go undetected/untreated and may result in disability.^32^

In a cross-sectional self-reported survey of early-stage lung cancer survivors^33^ only 23% met physical activity recommendations compared to 33% for the general US adult population.^34^ The mean (EQ-5D) health utility for lung cancer survivors who completed treatment >30 days previously is 0.79,^17,35^ much lower than the general US adult population (0.87)^36^ and those without chronic conditions (0.89).^36^ While suggesting unique opportunities to improve lung cancer health, these findings also imply cautionary considerations. Among them, lung cancer patients are somewhat different from other cancers due to higher median age at diagnosis, lifetime tobacco exposure, and prevalence of comorbidities.^8^ As well, curative-intent lung cancer treatment may have unique effects related to cardiopulmonary health since part of the treatment requires local destruction and/or removal of lung tissue which may impede cardiopulmonary function and exercise capacity.

Exercise interventions in lung cancer survivors commonly take place as part of comprehensive, multidisciplinary pulmonary rehabilitation programs, either in the preoperative or postoperative context.^37^ Recent systematic reviews suggest that preoperative exercise improves cardiopulmonary fitness^38,39^ and may reduce the risk of surgical complications,^40-43^ while postoperative exercise may improve functional and/or exercise capacity and QoL.^42-45^ While these findings provide supporting evidence for exercise in lung cancer survivorship, the authors of the systematic reviews raise concerns for volunteer and selection bias and inadequate sample size for the included studies, and note a lack of large randomized controlled trials (RCTs) of exercise interventions for lung cancer survivors.^40,43-45^ A more recent RCT^46^ examining the impact of physical activity/exercise on fatigue in advanced lung cancer survivors showed no difference compared to usual care; however, in exploratory analyses of 38 (34%) of participants who increased physical activity/exercise during the intervention period, there was a significant reduction in fatigue symptoms.^46^

Cancer rehabilitation has been suggested to be effective in improving function and QoL in cancer survivors and may be cost-effective in reducing healthcare costs.^32^ Exercise plays an important role in rehabilitation and may have role in cancer rehabilitation to improve physical and psychological outcomes, decrease treatment-related morbidity, increase treatment options, and potentially reduce costs.^32^ Such service may also assist lung cancer survivors overcome the physical and cardiopulmonary limitations, chronic disease management for COPD, heart failure, diabetes, and the associated psychological distress. Overcoming patient self-blame and stigma^47^ may be equally important to ensure adherence. Home-based interventions may also need to be incorporated to maintain exercise and/or facilitate program uptake for patients who have barriers for facility-based programs.

Our study has limitations. First, we assumed a health utility improvement reported in LIFE which enrolled older adults without cancer and, therefore, may not translate to the lung cancer population. However, there are several advantages in choosing LIFE including: the advanced age of participants; multicenter and US setting; and the availability of cost-effectiveness data^16^ which maximize accuracy and real-life implications. In addition, recent systematic reviews suggest that exercise training in lung cancer patients may produce benefits in improving physical fitness^42,43^ and, possibly, physical health QoL (in a meta-analysis of four studies involving 138 surgically-treated lung cancer patients^42^). Furthermore, the QoL benefits in LIFE extended across participants with and without comorbidity and among those with varying levels of fitness, suggesting that the LIFE exercise intervention could plausibly work among lung cancer patients. Second, the utility instrument used in LIFE (Quality of Well-being Scale) may have different psychometric properties including responsiveness to change compared to the EQ-5D. Nevertheless, similarities between these instruments include the use of preference-based scales (0-1) and the utility improvement in LIFE^16^ (0.05 QALYs over a median of 2.6 years) resembles the magnitude of EQ-5D utility improvement for COPD patients undergoing eight weeks of pulmonary rehabilitation,^48^ meeting the threshold for minimal clinically important difference in lung cancer.^49^ Third, we did not include possible negative health consequences related to exercise. However, in LIFE there was no difference in major adverse events related to exercise between the exercise and health education programs.^16^ In addition, the US Preventative Task Force suggests that potential harms are at most small and favors interventions to promote physical activity in individuals with^50^ and without^51^ cardiovascular disease risk factors. Fourth, we obtained DFS and OS data from patients undergoing lung cancer resection surgery and therefore may have limited generalizability to those undergoing definitive radiation or concurrent chemoradiation. However, the wide range of 5-year DFS and OS rates tested in our sensitivity analyses included stage IA patients undergoing lobectomy (respective rates ∼80% ^23^ and 60%^52^) and stage IIIA patients undergoing concurrent chemoradiation (respective rates ∼25% and 30%^53^) and therefore facilitated inferences of cost-effectiveness for a broad range of treatment modalities. Fifth, we did not incorporate possible benefits of exercise on functional changes associated with aging, chronic disease control, healthcare utilization including pulmonary rehabilitation, or survival; however, including these benefits would likely favor the cost-effectiveness of the LIFE exercise program. Finally, while this cost-effectiveness model strove to incorporate high-level evidence from a focused population of lung cancer patients, we lack a definitive RCT evaluating the LIFE exercise intervention in stage I-IIIA NSCLC patients. To identify model inputs for the CEA, we turned to research involving elderly non-cancer adults (for exercise program utility and cost), and breast cancer patients (for probability of exercise uptake). We measured the importance of these assumptions including a lower (40%) probability of increasing exercise and (0.01) QALY improvement related to exercise in our one-way sensitivity analyses, though the possibility of bias remains.

Our model did not include immune checkpoint blockade (ICB) costs since they are not currently standard therapy for stage III or earlier NSCLC, although ICB therapy is moving rapidly to earlier-stage disease. A recent RCT comparing ICB with anti-programmed death (PD) ligand 1 antibody durvalumab after platinum-based chemoradiotherapy in patients with unresectable stage III NSCLC found longer progression-free survival with ICB compared to placebo.^54^ In a subgroup analysis, patients who started durvalumab <14 days of completing chemoradiation had a progression-free survival hazard ratio (HR) of 0.39, compared to those patients who received ICB 14-56 days of completing chemoradiation who had an HR of 0.63. In addition to underlying comorbidities, these findings may support the hypothesis that ICB integrated more proximally, or potentially concurrently, with radiation may improve outcomes for patients with locally-advanced NSCLC treated with definitive chemoradiation. ANVIL is an ongoing NCI Cooperative Group RCT testing nivolumab (anti-PD-1 antibody) as adjuvant therapy after surgical resection in stage IB-IIIA (https://clinicaltrials.gov/ct2/show/NCT02595944), the results of which will further clarify the role of ICBs in patients with curable non-metastatic disease. While our model did not include the possible benefit of consolidation ICB therapy following or during traditional curative-intent treatment, including such benefit would only further improve the cost-effectiveness of exercise in our model. Furthermore, future studies examining the utility of exercise in lung cancer may include not only improving function and QoL, but also possible synergistic effects of exercise, radiation and ICB on lung cancer outcomes.

We conclude that in a simulation model, the LIFE exercise program is cost-effective compared to usual care in improving QoL in lung cancer survivors following curative-intent treatment from an organizational, but not societal perspective. Strategies to effectively increase exercise remotely may be more cost-effective than in-facility strategies for these patients.

## Abbreviations

CEA: cost-effectiveness analysis
COPD: chronic obstructive pulmonary disease
DFS: disease-free survival
EQ-5D: EuroQoL-5 Dimensions
HI: high intensity
HR: hazard ratio
ICB: immune checkpoint blockade
ICER: incremental cost-effectiveness ratio
LIFE: Lifestyle Interventions and Independence for Elders
LMI: low-moderate intensity
NSCLC: non-small cell lung cancer
OS: overall survival
PD: programmed death
QALY: quality-adjusted life-years
QoL: quality of life
RCT: randomized clinical trial
SD: standard deviation
SEER: Surveillance, Epidemiology, and End Results
UC: usual care
US: United States
WTP: willingness-to-pay

## References

1. Dinan MA, Curtis LH, Carpenter WR, et al. Stage migration, selection bias, and survival associated with the adoption of positron emission tomography among medicare beneficiaries with non-small-cell lung cancer, 1998-2003. J Clin Oncol. 2012;30(22):2725–2730.

2. Agostini P, Cieslik H, Rathinam S, et al. Postoperative pulmonary complications following thoracic surgery: Are there any modifiable risk factors? Thorax. 2010;65(9):815–818.

3. Ha D, Choi H, Zell K, et al. Association of impaired heart rate recovery with cardiopulmonary complications after lung cancer resection surgery. J Thorac Cardiovasc Surg. 2015;149(4):1168-73.e3.

4. Wang JS, Abboud RT, Wang LM. Effect of lung resection on exercise capacity and on carbon monoxide diffusing capacity during exercise. Chest. 2006;129(4):863–872.

5. Brunelli A, Xiume F, Refai M, et al. Evaluation of expiratory volume, diffusion capacity, and exercise tolerance following major lung resection: A prospective follow-up analysis. Chest. 2007;131(1):141–147.

6. Win T, Groves AM, Ritchie AJ, Wells FC, Cafferty F, Laroche CM. The effect of lung resection on pulmonary function and exercise capacity in lung cancer patients. Respir Care. 2007;52(6):720–726.

7. Carver JR, Shapiro CL, Ng A, et al. American society of clinical oncology clinical evidence review on the ongoing care of adult cancer survivors: Cardiac and pulmonary late effects. J Clin Oncol. 2007;25(25):3991–4008.

8. Islam KM, Jiang X, Anggondowati T, Lin G, Ganti AK. Comorbidity and survival in lung cancer patients. Cancer Epidemiol Biomarkers Prev. 2015;24(7):1079–1085.

9. Poghosyan H, Stock S, Kennedy Sheldon L, Cromwell J, Cooley ME, Nerenz DR. Racial disparities in health-related quality of life after lung cancer surgery: Findings from the cancer care outcomes research and surveillance consortium. J Thorac Oncol. 2015;10(10):1404–1412.

10. Denlinger CS, Ligibel JA, Are M, et al. Survivorship: Healthy lifestyles, version 2.2014. J Natl Compr Canc Netw. 2014;12(9):1222–1237.

11. Buffart LM, Kalter J, Sweegers MG, et al. Effects and moderators of exercise on quality of life and physical function in patients with cancer: An individual patient data meta-analysis of 34 RCTs. Cancer Treat Rev. 2017;52:91-104.

12. Schmitz KH, Courneya KS, Matthews C, et al. American college of sports medicine roundtable on exercise guidelines for cancer survivors. Med Sci Sports Exerc. 2010;42(7):1409– 1426.

13. Rock CL, Doyle C, Demark-Wahnefried W, et al. Nutrition and physical activity guidelines for cancer survivors. CA Cancer J Clin. 2012;62(4):243–274.

14. Pahor M, Guralnik JM, Ambrosius WT, et al. Effect of structured physical activity on prevention of major mobility disability in older adults: The LIFE study randomized clinical trial. JAMA. 2014;311(23):2387–2396.

15. National cancer institute: Surveillance, epidemiology, and end results program. cancer stats facts: Lung and bronchus cancer. https://seer.cancer.gov/statfacts/html/lungb.html. Accessed 10/16, 2017.

16. Groessl EJ, Kaplan RM, Castro Sweet CM, et al. Cost-effectiveness of the LIFE physical activity intervention for older adults at increased risk for mobility disability. J Gerontol A Biol Sci Med Sci. 2016;71(5):656–662.

17. Tramontano AC, Schrag DL, Malin JK, et al. Catalog and comparison of societal preferences (utilities) for lung cancer health states: Results from the cancer care outcomes research and surveillance (CanCORS) study. Med Decis Making. 2015;35(3):371–387.

18. Gordon LG, DiSipio T, Battistutta D, et al. Cost-effectiveness of a pragmatic exercise intervention for women with breast cancer: Results from a randomized controlled trial. Psychooncology. 2017;26(5):649–655.

19. Speck RM, Courneya KS, Masse LC, Duval S, Schmitz KH. An update of controlled physical activity trials in cancer survivors: A systematic review and meta-analysis. J Cancer Surviv. 2010;4(2):87–100.

20. Bradley CJ, Yabroff KR, Mariotto AB, Zeruto C, Tran Q, Warren JL. Antineoplastic treatment of advanced-stage non-small-cell lung cancer: Treatment, survival, and spending (2000 to 2011). J Clin Oncol. 2017:JCO2016694166.

21. Sanders GD, Neumann PJ, Basu A, et al. Recommendations for conduct, methodological practices, and reporting of cost-effectiveness analyses: Second panel on cost-effectiveness in health and medicine. JAMA. 2016;316(10):1093–1103.

22. Troiano RP, Berrigan D, Dodd KW, Masse LC, Tilert T, McDowell M. Physical activity in the united states measured by accelerometer. Med Sci Sports Exerc. 2008;40(1):181–188.

23. Dziedzic DA, Rudzinski P, Langfort R, Orlowski T, Polish Lung Cancer Study Group (PLCSG). Risk factors for local and distant recurrence after surgical treatment in patients with non-small-cell lung cancer. Clin Lung Cancer. 2016;17(5):e157–e167.

24. Social Security Actuarial life table. https://www.ssa.gov/oact/STATS/table4c6.html. Accessed 10/16, 2017.

25. Neumann PJ, Cohen JT, Weinstein MC. Updating cost-effectiveness--the curious resilience of the $50,000-per-QALY threshold. N Engl J Med. 2014;371(9):796–797.

26. van Waart H, van Dongen JM, van Harten WH, et al. Cost-utility and cost-effectiveness of physical exercise during adjuvant chemotherapy. Eur J Health Econ. 2017.

27. Kampshoff CS, van Dongen JM, van Mechelen W, et al. Long-term effectiveness and cost-effectiveness of high versus low-to-moderate intensity resistance and endurance exercise interventions among cancer survivors. J Cancer Surviv. 2018.

28. May AM, Bosch MJ, Velthuis MJ, et al. Cost-effectiveness analysis of an 18-week exercise programme for patients with breast and colon cancer undergoing adjuvant chemotherapy: The randomised PACT study. BMJ Open. 2017;7(3):e012187-2016-012187.

29. American cancer society: Economic impact of cancer. https://www.cancer.org/cancer/cancer-basics/economic-impact-of-cancer.html. Published 1/3/2018. Updated 2018. Accessed 02/26, 2018.

30. Guy GP,Jr, Yabroff KR, Ekwueme DU, Rim SH, Li R, Richardson LC. Economic burden of chronic conditions among survivors of cancer in the united states. J Clin Oncol. 2017;35(18):2053–2061.

31. Institute of medicine, committee on quality of healthcare in america. crossing the quality chasm: A new health system for the 21st century. washington, DC. National Academy Press. 2001.

32. Silver JK, Baima J, Mayer RS. Impairment-driven cancer rehabilitation: An essential component of quality care and survivorship. CA Cancer J Clin. 2013;63(5):295–317.

33. Krebs P, Coups EJ, Feinstein MB, et al. Health behaviors of early-stage non-small cell lung cancer survivors. J Cancer Surviv. 2012;6(1):37–44.

34. Evenson KR, Wen F, Herring AH. Associations of accelerometry-assessed and self-reported physical activity and sedentary behavior with all-cause and cardiovascular mortality among US adults. Am J Epidemiol. 2016;184(9):621–632.

35. Hung HY, Wu LM, Chen KP. Determinants of quality of life in lung cancer patients. J Nurs Scholarsh. 2018;50(3):257–264.

36. Jia H, Lubetkin EI. Estimating EuroQol EQ-5D scores from population healthy days data. Med Decis Making. 2008;28(4):491–499.

37. Bade BC, Thomas DD, Scott JB, Silvestri GA. Increasing physical activity and exercise in lung cancer: Reviewing safety, benefits, and application. J Thorac Oncol. 2015;10(6):861–871.

38. Loewen GM, Watson D, Kohman L, et al. Preoperative exercise Vo2 measurement for lung resection candidates: Results of cancer and leukemia group B protocol 9238. J Thorac Oncol. 2007;2(7):619–625.

39. Brunelli A, Kim AW, Berger KI, Addrizzo-Harris DJ. Physiologic evaluation of the patient with lung cancer being considered for resectional surgery: Diagnosis and management of lung cancer, 3rd ed: American college of chest physicians evidence-based clinical practice guidelines. Chest. 2013; 143(5 Suppl):e166S-e190S.

40. Cavalheri V, Granger C. Preoperative exercise training for patients with non-small cell lung cancer. Cochrane Database Syst Rev. 2017;6:CD012020.

41. Sebio Garcia R, Yanez Brage MI, Gimenez Moolhuyzen E, Granger CL, Denehy L. Functional and postoperative outcomes after preoperative exercise training in patients with lung cancer: A systematic review and meta-analysis. Interact Cardiovasc Thorac Surg. 2016;23(3):486–497.

42. Ni HJ, Pudasaini B, Yuan XT, Li HF, Shi L, Yuan P. Exercise training for patients pre- and postsurgically treated for non-small cell lung cancer: A systematic review and meta-analysis. Integr Cancer Ther. 2017;16(1):63–73.

43. Driessen EJ, Peeters ME, Bongers BC, et al. Effects of prehabilitation and rehabilitation including a home-based component on physical fitness, adherence, treatment tolerance, and recovery in patients with non-small cell lung cancer: A systematic review. Crit Rev Oncol Hematol. 2017;114:63-76.

44. Cavalheri V, Tahirah F, Nonoyama M, Jenkins S, Hill K. Exercise training undertaken by people within 12 months of lung resection for non-small cell lung cancer. Cochrane Database Syst Rev. 2013;(7):CD009955. doi(7):CD009955.

45. Cavalheri V, Tahirah F, Nonoyama M, Jenkins S, Hill K. Exercise training for people following lung resection for non-small cell lung cancer - a cochrane systematic review. Cancer Treat Rev. 2014;40(4):585–594.

46. Dhillon HM, Bell ML, van der Ploeg HP, et al. Impact of physical activity on fatigue and quality of life in people with advanced lung cancer: A randomized controlled trial. Ann Oncol. 2017;28(8):1889–1897.

47. Scott N, Crane M, Lafontaine M, Seale H, Currow D. Stigma as a barrier to diagnosis of lung cancer: Patient and general practitioner perspectives. Prim Health Care Res Dev. 2015;16(6):618–622.

48. Nolan CM, Longworth L, Lord J, et al. The EQ-5D-5L health status questionnaire in COPD: Validity, responsiveness and minimum important difference. Thorax. 2016;71(6):493–500.

49. Pickard AS, Neary MP, Cella D. Estimation of minimally important differences in EQ-5D utility and VAS scores in cancer. Health Qual Life Outcomes. 2007;5:70-7525-5-70.

50. LeFevre ML, U.S. Preventive Services Task Force. Behavioral counseling to promote a healthful diet and physical activity for cardiovascular disease prevention in adults with cardiovascular risk factors: U.S. preventive services task force recommendation statement. Ann Intern Med. 2014;161(8):587–593.

51. US Preventive Services Task Force, Grossman DC, Bibbins-Domingo K, et al. Behavioral counseling to promote a healthful diet and physical activity for cardiovascular disease prevention in adults without cardiovascular risk factors: US preventive services task force recommendation statement. JAMA. 2017;318(2):167–174.

52. van den Berg LL, Klinkenberg TJ, Groen HJ, Widder J. Patterns of recurrence and survival after surgery or stereotactic radiotherapy for early stage NSCLC. J Thorac Oncol. 2015;10(5):826–831.

53. Senan S, Brade A, Wang LH, et al. PROCLAIM: Randomized phase III trial of pemetrexed-cisplatin or etoposide-cisplatin plus thoracic radiation therapy followed by consolidation chemotherapy in locally advanced nonsquamous non-small-cell lung cancer. J Clin Oncol. 2016;34(9):953–962.

54. Sharabi AB, Lim M, DeWeese TL, Drake CG. Radiation and checkpoint blockade immunotherapy: Radiosensitisation and potential mechanisms of synergy. Lancet Oncol. 2015;16(13):e498–509.

